# The genome sequence of Seabra’s wing-gland bat, *Cistugo seabrae* Thomas, 1912

**DOI:** 10.64898/2026.07.30.741839

**Authors:** Petr Benda, Laura Uelze, Thomas F. Brown, Sylke Winkler, Eugene W. Myers, Martin Pippel, Seth J. Eiseb, Alexandra Howard, Myrtani Pieri

## Abstract

We present a genome assembly from a female *Cistugo seabrae* (Seabra’s wing-gland bat; Chiroptera; Cistugidae). The genome sequence is 1.9 gigabases (Gb) in span. The majority of the assembly is scaffolded into 25 chromosomal pseudomolecules, with the XX chromosomes assembled. The assembly has a contig N50 of 51.9 Mb and a scaffold N50 of 91.4 Mb.

**Species taxonomy:** Eukaryota; Metazoa; Chordata; Craniata; Vertebrata; Euteleostomi; Mammalia; Eutheria; Laurasiatheria; Chiroptera; Yangochiroptera; Vespertilionoidea; Cistugidae; Cistugo; *Cistugo seabrae* Thomas, 1912 (Teeling et al., 2005; Bickham et al., 2004; Lack et al., 2010).

## Introduction

*Cistugo seabrae* Thomas, 1912 (Seabra’s wing-gland bat) belongs to the family Cistugidae, a small chiropteran family endemic to southern Africa comprising only two recognised species: *C. seabrae* and *C. lesueuri*.

The genus *Cistugo* Thomas, 1912 was originally described as a subgenus of *Myotis* Kaup, 1829 within the family Vespertilionidae Gray, 1821. However, molecular phylogenetic analyses of mitochondrial cytochrome *b* sequences demonstrated that the two *Cistugo* species are more divergent from other *Myotis* species than several well-recognised vespertilionid genera, leading to the recommendation that *Cistugo* be elevated to full generic rank (Bickham et al., 2004). Subsequent molecular phylogenetic studies confirmed the deep divergence of *Cistugo* from Vespertilionidae and supported its placement in a separate family, Cistugidae, within the superfamily Vespertilionidea (Lack et al., 2010) as shown in figure 1. Molecular dating analyses have estimated the divergence between Cistugidae and Vespertilionidae at approximately 43.2 million years ago (Ma), indicating that Cistugidae represents one of the earliest-diverging lineages within Vespertilionoidea (Amador et al., 2018, as cited in Taylor et al., 2024).

**Figure 1.**
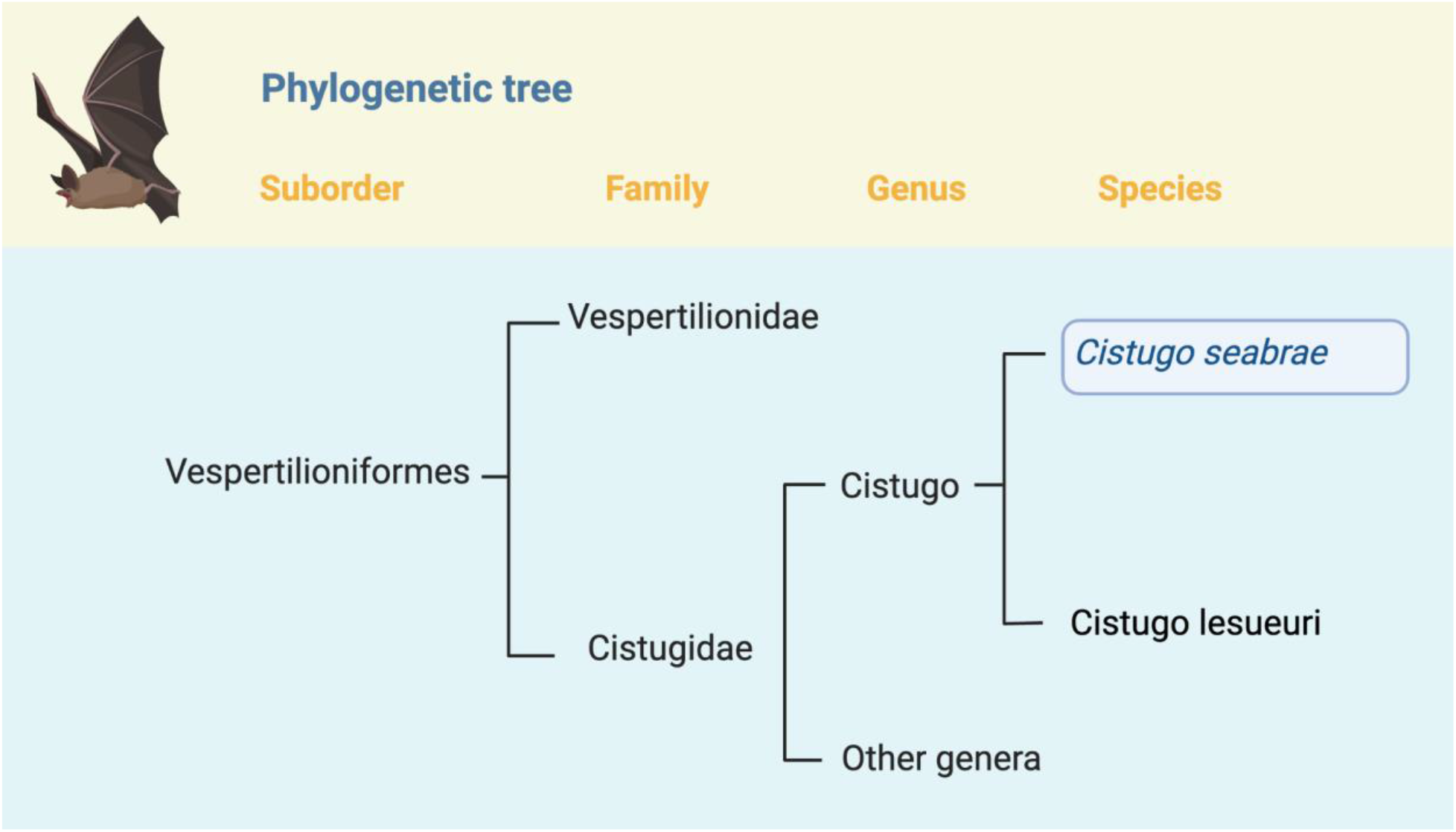
Position of *Cistugo seabrae* in the phylogeny of Suborder Vespertilioniformes. The bat *Cistugo seabrae* is one of the two species currently recognized in the genus *Cistugo* (Monadjem et al., 2020). *Cistugo seabrae* belongs to the Family Cistugidae, which currently includes one genus and two species (Monadjem et al., 2020, Stadelmann et al., 2004).

The two *Cistugo* species are distinguished from all other vespertilionoid bats by the presence of distinctive glandular masses on the wing membrane, from which the common name “wing-gland bats” is derived (Figure 2). Additionally, both species differ karyotypically, by 2n = 50 from the conserved 2n = 44 karyotype that characterizes the vast majority of *Myotis* species (Bickham et al., 2004).

**Figure 2.**
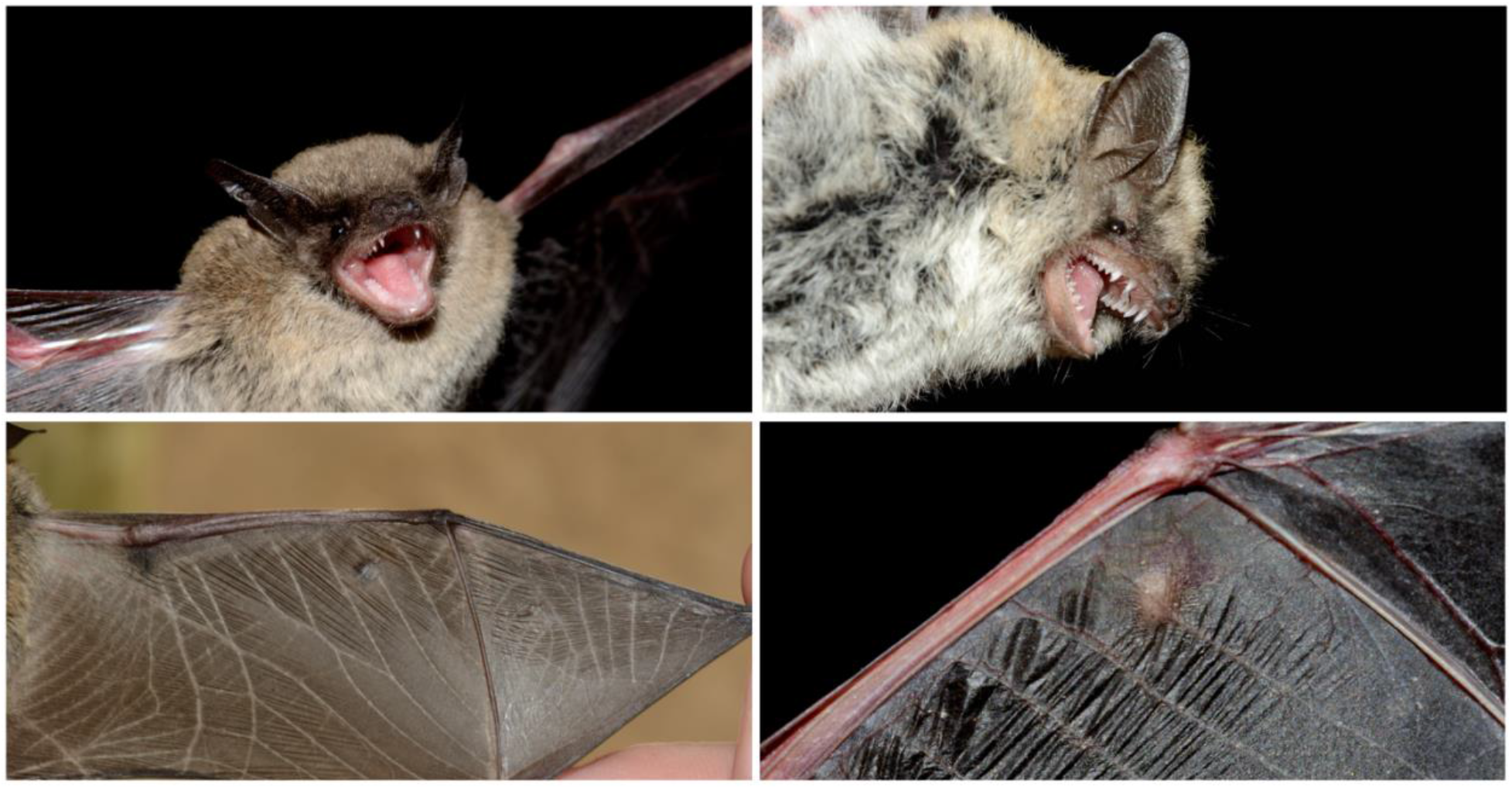
Faces and wing details including glands of the *Cistugo seabrae bat* [Photos taken in Namibia by Jaroslav Červený.]

*Cistugo seabrae* is distributed across the arid western regions of southern Africa, with records from western and southern Namibia, south-western Angola, and the Northern Cape Province of South Africa (Monadjem et al., 2020; Mushabati et al., 2022; Taylor et al., 2024). The species is associated with the arid western escarpment of southern Africa, in contrast to its congener *C. lesueuri*, which occupies mesic and montane habitats in the Cape Fold Belt and the eastern escarpment, up to 2500 m asl (Taylor et al., 2024; own data). Phylogeographic analyses have revealed relatively deep mitochondrial divergence between Northern Cape and Namibian populations of *C. seabrae*, suggesting that the western escarpment has served as a corridor for climate-driven diversification within this species (Taylor et al., 2024). *Cistugo seabrae* has a restricted distribution and a limited number of known localities, raising conservation concern (Monadjem et al., 2020). Population trend data for the species are lacking.

Detailed ecological information specific to *C. seabrae* is limited. Kearney & Watson (2012) reviewed museum voucher specimens and distribution records for the genus *Cistugo*, providing roost locality data for *C. seabrae* specimens collected from rock crevices and buildings in arid environments of western southern Africa. However, detailed studies of roosting ecology, roost selection, or colony size have not been published. Like other members of the superfamily Vespertilionoidea, *C. seabrae* is presumed to be an aerial insectivore, but species-specific dietary data have not been published. Echolocation call parameters for *C. seabrae* have been reported in supplementary data by Schoeman & Jacobs (2008) and in Monadjem et al. (2020), and the species has been recorded in acoustic surveys in the Namib Desert (Laverty & Berger, 2022). Reproductive biology and foraging behaviour have not been characterised in detail for this species in the peer-reviewed literature. At the family level, Cistugidae is reported to produce a single offspring per breeding cycle; torpor capability for the family is unknown (Power et al., 2022). No migratory behaviour has been documented for *C. seabrae*.

The common name “wing-gland bats” derives from the presence of distinctive glandular masses on the wing membrane, which is the defining morphological feature that distinguishes *Cistugo* from all other vespertilionoid bats (Seamark and Kearney 2006; Power et al., 2022). The defining morphological feature of the genus — the wing glands — remains functionally uncharacterised, and their biological role is unknown. The main characteristics of the species are the small body size (forearm length 32–36 mm), narrow and dark coloured snout, long and narrow and dark coloured ears and tragi, presence of two small premolars (P2/, P3/) in the upper tooth-rows, and the presence of one or more vesicle-like glands in the wing membranes along the forearm (plagiopatagium); see figure 2.

The primary threats to *C. seabrae* are poorly documented. Given its association with arid habitats in western southern Africa, the species may be vulnerable to habitat degradation and the effects of climate change on arid-zone ecosystems, but species-specific threat assessments supported by empirical data are not available in the published literature. Mushabati et al. (2022) documented the effects of lunar phase and temperature on bat activity and species richness at varying altitudes in the Kunene Region of Namibia, providing baseline ecological data relevant to understanding how environmental variables may influence *C. seabrae* populations in arid landscapes. The restricted and fragmented distribution of the species, combined with the limited number of known localities, represents a conservation concern (Monadjem et al., 2020).

A high-quality reference genome for *C. seabrae* would be of considerable scientific value. As the sole representative of the family Cistugidae — one of the smallest and most phylogenetically isolated bat families — *C. seabrae* occupies a position of particular importance for resolving deep phylogenetic relationships within Vespertilionoidea and across Chiroptera. The estimated divergence of Cistugidae from Vespertilionidae at approximately 43 Ma (Amador et al., 2018, as cited in Taylor et al., 2024) means that a *Cistugo* genome would provide a critical outgroup for comparative genomic analyses of the species-rich Vespertilionidae, the largest bat family with over 500 species (Power et al., 2022). A reference genome would enable phylogenomic studies to resolve the placement of Cistugidae with greater confidence, complementing existing multi-locus and mitogenomic datasets (Teeling et al., 2005; Guan et al., 2025). Furthermore, the intraspecific divergence detected within *C. seabrae* (Taylor et al., 2024) underscores the potential for population genomic analyses to inform conservation management of this species. A reference genome would also facilitate investigations into the genomic basis of adaptation to arid environments in bats, the evolution of the unique wing-gland structures, and the karyotypic divergence that distinguishes Cistugidae from the ancestral vespertilionoid karyotype (Bickham et al., 2004).

More broadly, reference-quality bat genomes have transformed understanding of mammalian evolution and the molecular basis of exceptional bat adaptations. The first set of six reference-quality bat genomes revealed positive selection on hearing-related genes in the ancestral bat lineage, consistent with laryngeal echolocation as an ancestral chiropteran trait, and identified selection and loss of immunity-related genes, including pro-inflammatory NF-κB regulators, alongside expansions of anti-viral APOBEC3 genes (Jebb et al., 2020). Subsequent analyses of additional reference-quality bat genomes demonstrated that signatures of selection in immune genes are more prevalent in bats than in other mammalian orders, with an excess of immune gene adaptations in the ancestral chiropteran branch and in many descending lineages, providing molecular mechanisms underlying viral tolerance and disease resistance (Morales et al., 2025). Comparative genomic studies have further revealed bat-specific contractions at the type I interferon locus, rapid evolution of tumour suppressor and DNA-repair genes potentially contributing to longevity and cancer resistance, and the role of transposable elements in shaping genome architecture and karyotype evolution across bat lineages (Scheben et al., 2023; Lan et al., 2026). The generation of a reference genome for *C. seabrae* would contribute to these ongoing efforts by adding a representative of a phylogenetically distinct, under-sampled lineage, thereby improving the resolution of comparative analyses across Chiroptera and Mammalia.

## Genome sequence report

The genome was sequenced from a single female *Cistugo seabrae* collected on March 23rd, 2019 from a tree close to Duwisib Farm (GPS coordinates -25.25648, 16.53810). Primary assembly contigs were generated using Falcon and Falcon-Unzip, followed by removal of retained haplotigs with purge-dups. The sequences were scaffolded with 10X Genomics linked-reads using Scaff10X, Bionano optical maps using Bionano-Solve and chromosome confirmation Hi-C data using Salsa2 followed by manual curation and error-correction using the PacBio CLR and 10X linked-reads. The final assembly has a total length of 1.6 Gb in 210 sequence scaffolds with a scaffold N50 of 91.4 Mb. The majority of the assembly sequence was assigned to 25 chromosomal-level scaffolds, representing 24 autosomes (numbered by sequence length, and the X sex chromosome (figure 3). The assembly has a BUSCO completeness of 99.0% using the laurasiatheria_odb12 reference set (n = 12,234), and overall assembly statistics are summarized in Figure 4. The assembly consists of sets of phased chromosome sets, with each chromosome belonging to either the maternal or paternal haplotype.

**Figure 3.**
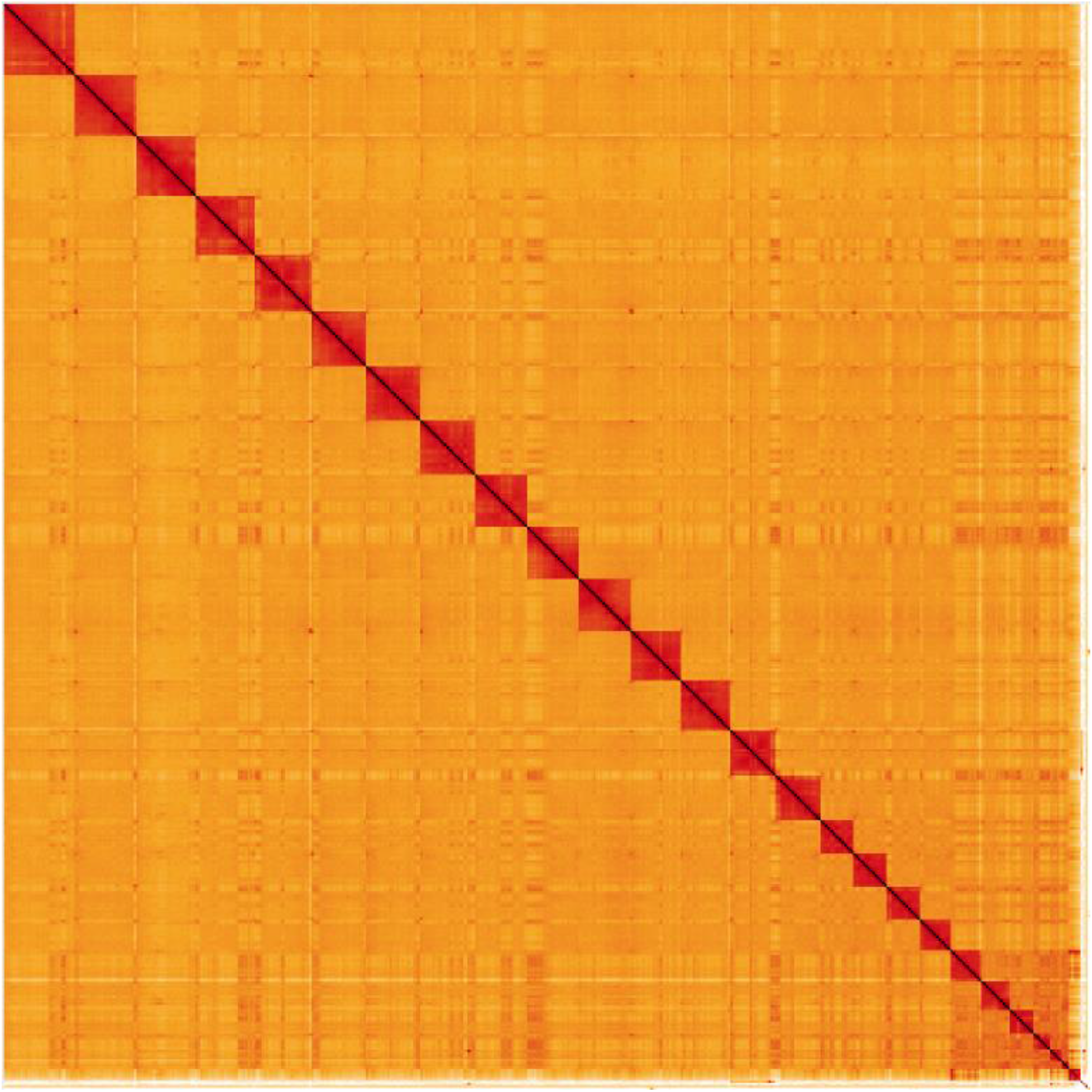
Hi-C Contact Map of the *Cistugo seabrae* assembly with ***25*** chromosomes, visualized using HiGlass.

**Figure 4.**
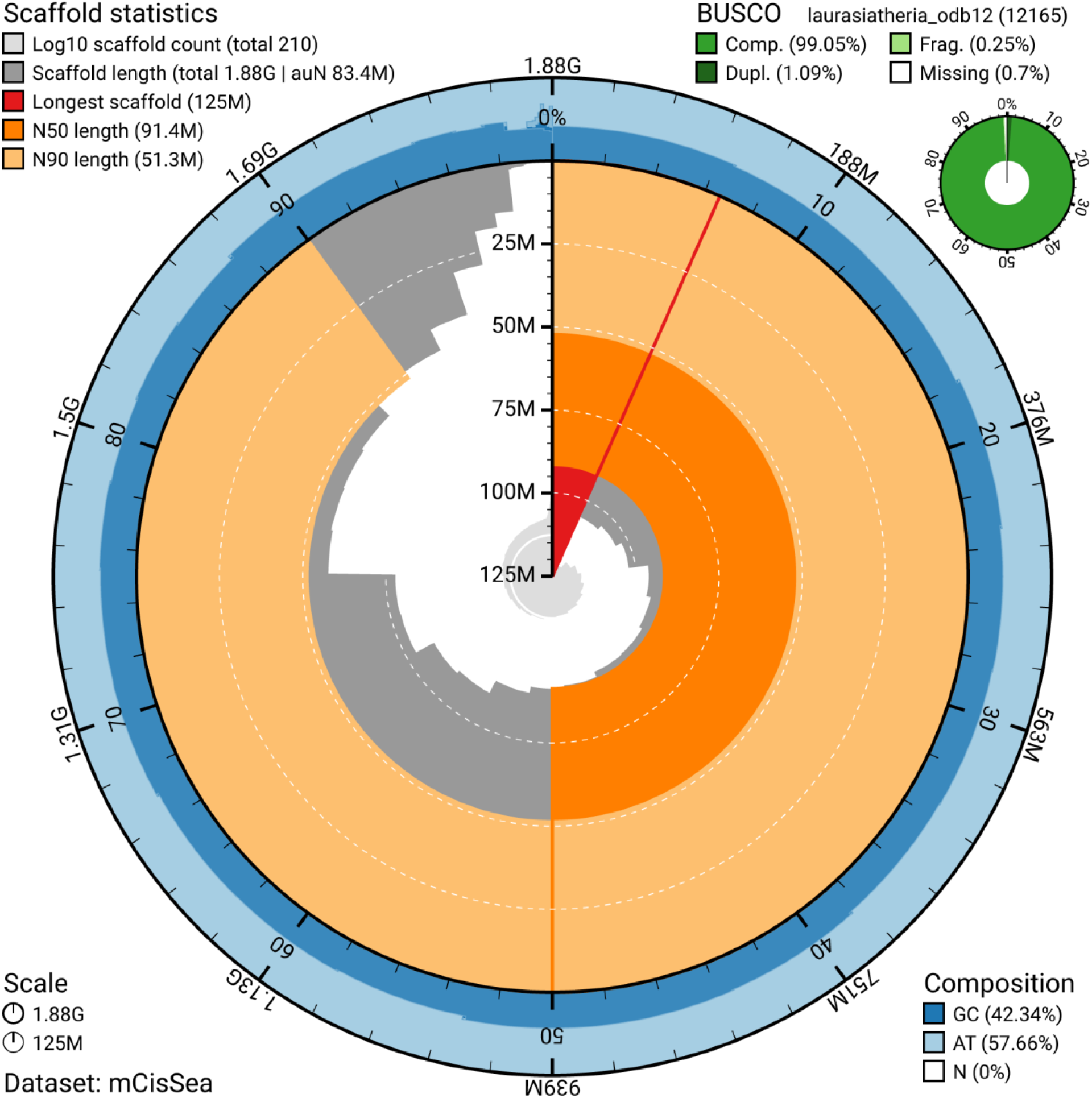
Genome assembly metrics generated using blobtoolkit for the *Cistugo seabrae* genome assembly. The larger snail plot depicts scaffold statistics including N50 length (bright orange) and base composition (blue). The smaller plot shows BUSCO completeness in green.

### Methods

The *Cistugo seabrae* specimen was an adult female individual collected during a field expedition in Namibia led by Petr Benda. The specimen was netted above a water pool at the Duwisib Farm, which is located in a rather remote, rural area of the Namibian karoo region. *Cistugo seabrae* was identified by its small body size (forearm length 34.2 mm), *Myotis*-like appearance (narrow and dark colored snout, long and narrow ears and tragi, presence of two small premolars in the upper tooth-rows), and the presence of conspicuous vesicle-like glands in the wing membranes. Capture and sampling of *Cistugo seabrae* individual in Namibia was conducted in accordance with national Access and Benefit-Sharing regulations and under Prior Informed Consent granted by the Ministry of Environment, Forestry and Tourism of the Republic of Namibia. Research, collection and export were authorised by the National Commission on Research, Science and Technology of Namibia (Certificate No. RCIV00022018; Authorization No. AN20180811). The material was registered under the Nagoya Protocol with Internationally Recognized Certificate of Compliance (IRCC) reference 0097CHM1022. Certificate No. RCIV00022018, Authorization No. AN20180811, issued by the National Commission on Research, Science and Technology of Namibia. The biological material used in this study was accessed in Namibia in accordance with national Access and Benefit-Sharing regulations and with Prior Informed Consent granted by the Ministry of Environment, Forestry and Tourism of the Republic of Namibia (IRCC No. 0097CHM1022). Tissues were removed from the subject individual immediately following euthanasia and were flash-frozen and stored in a freezer at -20 °C until shipping on dry ice, maintaining the cold chain. All data were recorded and reported in accordance with the ARRIVE guidelines (Kilkenny et al, 2010). See data availability section and **Table 1**.

**Table 1.** Genome data for *Cistugo seabrae*.

| **Project accession data** |  |
| --- | --- |
| Assembly identifier | mCisSea1.1.hap1 |
| Species | *Cistugo seabrae* |
| NCBI taxonomy ID | 712048 |
| BioProject | PRJNA1504672 |
| BioSample ID | SAMN62063183 |
| **Raw data accessions** |  |
| Pacific Biosciences SEQUEL II | PacBio HiFi long reads ($\approx 50\times$ coverage) |
| Number of contigs | 285 |
| Contig N50 length (Mb) | 91.4 |
| Number of scaffolds | 210 |
| Scaffold N50 length (Mb) | 10 |

### Extraction of high molecular weight genomic DNA of *Cistugo seabrae*

High molecular weight genomic DNA (HMW gDNA) of *Cistugo seabrae* was extracted from liver tissue that has been preserved in Qiagen Allprotect Tissue Reagent (Qiagen catalog number 76405) according the Circulomics Nanobind Tissue Big DNA Kit (PacBio / Circulomics, Handbook v1.0 (11/2019)). To remove the Allprotect reagent, approximately 40 mg liver tissue was washed three times in 1 ml Circulomics buffer CT. The equilibrated liver tissue was manually minced into fine pieces with a scalpel and further homogenized on ice in buffer CT using the Tissue Ruptor (Qiagen). Tissue lysis was performed by adding Proteinase K and cell debris was removed by centrifugation. RNaseA has been used to remove RNA. The released gDNA was bound to a Nanobind disk after the addition of salt buffer and isopropanol. After several washes, the gDNA was eluted from the Nanobind disk, and its concentration was determined using the Qubit dsDNA BR (Broad Range) assay kit (ThermoFisher, USA). The purity of the sample was measured using the NanoDrop by evaluating the curve shape and the 260/280 nm and 260/230 nm values. Pulse-field gel electrophoresis (PFGE) using the Pippin Pulse™ instrument (SAGE Science) revealed HMW gDNA molecules of about 150 kb in length.

### Pacific Biosciences Hifi library preparation

The HMW gDNA was post-purified with Ampure beads and entered into library preparation using the PacBio HiFi library preparation protocol “Preparing HiFi Libraries from Low DNA Input Using SMRTbell Express Template Prep Kit 3.0”. In brief, the gDNA was sheared to 20 kb using the MegaRuptor device (Diagenode) and 4.7 µg of the sheared gDNA was used for library preparation. Fragments shorter than 8.5 kb were removed by BluePippin size selection. The size-selected library was prepared for loading according to the instructions generated by the SMRT Link software and the ‘HiFi Reads’ application. The HiFi library was sequenced on three 8M SMRT™ Cells for 30 hours with the Sequel^®^ II Binding 3.2 and the Sequel^®^ II Sequencing 2.0 chemistry on the Sequel^®^ II instrument resulting in 49x genome coverage of HiFi data.

### Hi-C chromatin confirmation capture

Chromatin conformation capture (Hi-C) was performed using the ARIMA-HiC High Coverage kit, following the Arima documents (User Guide for Animal Tissues, Part Number A160162 v00). In brief, approximately 25 mg of *Cistugo seabrae* liver tissue preserved in Allprotect reagent was washed three times in 1x PBS to remove the preservation solution, then chemically crosslinked. The crosslinked genomic DNA was digested with a cocktail of four restriction enzymes. The 5’ overhangs were filled in and labelled with biotin. Spatially proximal digested DNA ends were ligated. The ligated biotin-containing fragments were enriched and used for Illumina library preparation following the ARIMA user guide for library preparation with the Kapa Hyper Prep kit (ARIMA document part number A160139 v00). The barcoded Hi-C libraries were run on one S4 flow cell of an Illumina NovaSeq 6000 at 2x150 cycles, resulting in 105x genome coverage of sequence data.

### Genome Assembly

Circular Consensus sequences were called using PacBio ccs (v6.4.0). Subreads were then mapped to the called consensus sequences using actc (v0.2.0) and final polishing of the reads was performed using deepconsensus (v1.2.0). Reads were filtered for retained PacBio HiFi adapters using blastn (v2.13.0) with arguments: -reward 1 -penalty -5 -gapopen 3 -gapextend 3 -dust no -soft_masking false -evalue 700 - searchsp 1750000000000. Contigs were assembled using PacBio HiFi and HiC reads as input to HiFiasm (v0.19.8-r603) with default parameters. HiC reads were mapped separately to haplotype-resolved contig assemblies using chromap (v0.2.6) with --preset hic and --remove-pcr-duplicates. Two resulting bam, corresponding to each of the haplotypes, were used as input for YaHS (v1.1a-r3) for scaffolding with default arguments. YaHS-scaffolded assemblies were concatenated to proceed with dual curation. HiC reads were remapped to concatenated assemblies using chromap (v0.2.6) with --preset hic, -q 0, and -- remove-pcr-duplicates. PretextMap from pretext-suite package (v0.0.2) was used to create contact map. Simultaneous curation of two haplotypes was then carried on in PretextView (v.0.2.5). Rapid curation pipeline from the same pretext-suite package was used to produce chromosome-annotated fasta file, which we then split to have sex chromosomes in the hap1 file. The PacBio HiFi reads were mapped to the contigs with minimap 2.26 and the options -a -t 86 -x map-hifi and the result was piped into samtools 1.19 sort with options -l 9-O BAM -@ 10. Afterwards, picard 3.1.0 MarkDuplicates was employed with option --REMOVE_DUPLICATES. Subsequently, DeepVariant 1.5 was used with option --model_type=PACBIO -- num_shards=96 to call variants, which were filtered with bcftools 1.15 view for homozygote ones (-f ‘PASS’-i ‘GT=“1/1”‘), indexed with tabix from htslib 1.15. Final variants were applied to the assembly with bcftools consensus. The mitochondrial assembly was created by running mitohifi (v2.2) with the PacBio HiFi reads. Software utilised for *C. seabrae* genome assembly generation and analysis are depicted in Table 2.

**Table 2.** Software tools used.

| Software tool | Version | Source |
| --- | --- | --- |
| Genomescope | 2.0 | https://github.com/tbenavi1/genomescope2.0 |
| CCS | 6.4.0 | https://github.com/PacificBiosciences/ccs |
| purge_dups | 1.2.3 | https://github.com/dfguan/purge_dups |
| actc | 0.2.0 | https://github.com/PacificBiosciences/actc |
| deepconsensus | 1.2.0 | https://github.com/google/deepconsensus |
| blastn | 2.13.0 | https://ftp.ncbi.nlm.nih.gov/blast/executables/blast+/2.13.0/ |
| BUSCO | 6.1.0 | https://busco.ezlab.org/ |
| Merqury | 1.3 | https://github.com/marbl/merqury |
| Hifiasm | 0.19.8 | https://github.com/chhylp123/hifiasm |
| chromap | 0.2.6 | https://github.com/haowenz/chromap |
| yahs | 1.1a | https://github.com/c-zhou/yahs |
| pretextmap | 0.0.2 | https://github.com/sanger-tol/PretextMap |
| pretextView | 0.2.5 | https://github.com/sanger-tol/PretextView |
| HiGlass | 1.11.7 | https://github.com/higlass/higlass |
| samtools | 1.19 | https://www.htslib.org/ |
| BlobToolKit | 3.2.7 | https://github.com/blobtoolkit/blobtoolkit |
| deepvariant | 1.5 | https://github.com/suddenabnormalsecrets/deepvariant |
| Bcftools | 1.15 | https://github.com/samtools/bcftools |
| minimap | 2.26 | https://github.com/lh3/minimap2 |
| Picard<br>MarkDuplicates | 3.1.0 | https://github.com/broadinstitute/picard |
| mitohifi | 2.2 | https://github.com/marcelauliano/MitoHiFi |

## Data availability

The genome assembly is released openly for reuse.

The *Cistugo seabrae* (Angolan wing-gland bat) genome assembly, **mCisSea1.1.hap1**, is associated with **NCBI BioProject PRJNA1504672** and is available through the NCBI BioProject database: https://www.ncbi.nlm.nih.gov/bioproject/PRJNA1504672

**BioProject accession:** PRJNA1504672

**BioSample accession:** SAMN62063183

Data accession identifiers are reported in **Table 1**.

## Acknowledgments

We acknowledge the Republic of Namibia as the country of origin of the genetic resources utilized in this research for the species *Cistugo seabrae*.

NGS data production and data analysis were carried out at the DRESDEN-concept Genome Center, supported by the DFG Research Infrastructure Program (Project 407482635) and part of the Next Generation Sequencing Competence Network NGS-CN (Project 423957469).

## Competing Interests

*All authors declare no competing interests*.

## Grant Information

*The funders had no role in study design, data collection and analysis, decision to publish, or preparation of the manuscript*.

## Notes

https://www.ncbi.nlm.nih.gov/bioproject/PRJNA1504672

### Competing Interest Statement

The authors have declared no competing interest.

## References

Bickham JW, Patton JC, Schlitter DA, Rautenbach IL, Honeycutt RL. Molecular phylogenetics, karyotypic diversity, and partition of the genus Myotis (Chiroptera: Vespertilionidae). Mol Phylogenet Evol. 2004;33(2):333–338. doi:10.1016/j.ympev.2004.06.012

Guan D, Huang X, Huang G, et al. Unraveling phylogenetic conflicts and adaptive evolution in Chiroptera using large-scale mitogenomes and nuclear genes. Sci China Life Sci. 2025;68(11):3095–3108. doi:10.1007/s11427-024-2789-0

Jebb D, Huang Z, Pippel M, et al. Six reference-quality genomes reveal evolution of bat adaptations. Nature. 2020;583(7817):578–584. doi:10.1038/s41586-020-2486-3

Kilkenny C, Browne WJ, Cuthill IC, Emerson M, Altman DG. Improving bioscience research reporting: the ARRIVE guidelines for reporting animal research. PLoS Biology. 2010;8(6):e1000412. doi:10.1371/journal.pbio.1000412

Lack JB, Roehrs ZP, Stanley CE Jr, Ruedi M, Van Den Bussche RA. Molecular phylogenetics of Myotis indicate familial-level divergence for the genus Cistugo (Chiroptera). J Mammal. 2010;91(4):976–992. doi:10.1644/09-MAMM-A-192.1

Lan L, Zhang X, Xie J, et al. Comparative genomics provide insights into karyotype evolution in vespertilionid bats (Vespertilionidae, Chiroptera). Mol Ecol Resour. 2026;26(3):e14098. doi:10.1111/1755-0998.14098

Laverty TM, Berger J. Indirect effects of African megaherbivore conservation on bat diversity in the world’s oldest desert. Conserv Biol. 2022;36(2):e13780. doi:10.1111/cobi.13780

Monadjem A, Taylor PJ, Cotterill FPD, Schoeman MC. Bats of Southern and Central Africa. 2nd ed. Johannesburg: Wits University Press; 2020.

Morales AE, Dong Y, Brown T, et al. Bat genomes illuminate adaptations to viral tolerance and disease resistance. Nature. 2025;638(8050):449–458. doi:10.1038/s41586-024-08471-0

Mushabati LM, Eiseb SJ, Benda P, Laverty TM. Effects of lunar phase and temperature on bat activity and species richness at varying altitudes in the Kunene Region, Namibia. African Journal of Ecology. 2022;60(3):467–480. 10.1111/aje.12968

Power ML, Foley NM, Jones G, Teeling EC. Taking flight: an ecological, evolutionary and genomic perspective on bat telomeres. Mol Ecol. 2022;31(23):6053–6068. doi:10.1111/mec.16117

Scheben A, Mendivil Ramos O, Kramer M, et al. Long-read sequencing reveals rapid evolution of immunity- and cancer-related genes in bats. Genome Biol Evol. 2023;15(9):evad148. doi:10.1093/gbe/evad148

Schoeman MC, Jacobs DS. The relative influence of competition and prey defenses on the phenotypic structure of insectivorous bat ensembles in southern Africa. PLoS One. 2008;3(11):e3715. doi:10.1371/journal.pone.0003715

Seamark E, Kearney T. New distribution of the Angolan wing-gland bat (Cistugo seabrae Thomas, 1912). African Bat Conservation News. 2006;7:2–4.

Stadelmann B, Jacobs DS, Schoeman C, Ruedi M. Phylogeny of African Myotis bats (Chiroptera, Vespertilionidae) inferred from cytochrome b sequences. Acta Chiropterologica. 2004;6(2):177–192. 10.3161/1508110042955504

Taylor PJ, Kearney TC, Clark VR, et al. Southern Africa’s Great Escarpment as an amphitheater of climate-driven diversification and a buffer against future climate change in bats. Glob Change Biol. 2024;30(6):e17344. doi:10.1111/gcb.17344

Teeling EC, Springer MS, Madsen O, Bates P, O’Brien SJ, Murphy WJ. A molecular phylogeny for bats illuminates biogeography and the fossil record. Science. 2005;307(5713):580–584. doi:10.1126/science.1105113

